# Mechanosensitive recruitment of BAF to the nuclear membrane inhibits nuclear E2F1 and Yap levels

**DOI:** 10.1101/803932

**Authors:** C.P. Unnikannan, Adriana Reuveny, Devora Tamar Grunberg, Talila Volk

## Abstract

Mechanotransduction has been implicated as an important factor in regulating cell cycle progression; however, the underlying mechanism has not been fully elucidated. Here, we describe a novel mechano-sensitive component, namely *barrier to autointegration factor*, (BAF), which regulates DNA endocycling in *Drosophila* muscle fibers. We show that BAF negatively regulates DNA endoreplication by inhibiting of the nuclear entrance of E2F1 and Yap/Yorkie, two key components in cell cycle control. Furthermore, BAF localization at the nuclear membrane is mechanosensitive, as it was downregulated in LINC mutant larval muscles, or following nuclear deformation caused by disruption of nucleus-sarcomere connections. BAF forms a protein complex with E2F1, which is sensitive to BAF phosphorylation. Knockdown of BAF kinase VRK1/Ball disrupted localization of BAF at the nuclear membrane and resulted in increased E2F1 nuclear levels. Taken together, our results reveal a novel mechanosensitive pathway controlling BAF phosphorylation and localization at the nuclear membrane, which in turn, represses nuclear accumulation of positive cell cycle regulators.

## Introduction

Mechanical signals that are transmitted across the nuclear membrane have been implicated in the regulation of cell cycle, epigenetic events, gene expression and other processes (Cho et al., 2017; Kirby and Lammerding, 2018). Although the molecular mechanism that responds to mechanical signals and further transmitting them into the nucleus is still elusive, nuclear translocation of specific transcription factors controlling cell proliferation, including Yes Associated Protein (YAP)/Yorkie as well as Megakarioblastic Leukemia 1 (MKL1), and Serum Response Factor (SRF), has been implicated as part of the mechanotransduction pathway (Aragona et al., 2013; Driscoll et al., 2015; Dupont et al., 2011; Ho et al., 2013; Kassianidou et al., 2019). Moreover, translocation of YAP into the nucleus has been linked to mechanically dependent opening of the nuclear pore complex (Elosegui-Artola et al., 2017). Notably, additional molecular information linking nuclear translocation of transcription factors with nuclear mechanical inputs is still missing.

The linker of nucleoskeleton and cytoskeleton (LINC) complex is a potential mediator of the mechanically induced nuclear entry of essential factors (Driscoll et al., 2015; Horn, 2014; Osmanagic-Myers et al., 2015). The LINC complex physically connects the cytoskeleton and the nucleoskeleton at the interface of the nuclear membrane (Horn, 2014; Mejat and Misteli, 2010; Mellad et al., 2010). It is composed of Nesprin family protein members, which associate on one end with distinct cytoskeletal components, and on the other, with SUN domain proteins at the perinuclear space (Rajgor and Shanahan, 2013; Zhang et al., 2002; Zhang et al., 2001). SUN domain proteins bind to the nuclear lamina components, thereby producing a physical link between the cytoskeleton and the nucleoskeleton (Cain and Starr, 2015; Lombardi et al., 2011; Padmakumar et al., 2005; Sosa et al., 2012). Recent results have indicated that in *Drosophila* larval muscles, the LINC complex is essential for arresting cell cycle progression in the muscle nuclei (myonuclei). Furthermore, this study showed that in the myonuclei of LINC mutant larvae the DNA undergoes additional rounds of DNA replication known as endoreplication, leading to elevated polyploidy (Brayson et al., 2018; Volk, 2012; Wang et al., 2018). These observations implicate the involvement of the LINC complex in the arrest of cell cycle progression in myonuclei of wild-type larvae. The molecular nature of the mechanism transducing the arrest of cell cycle progression by the LINC complex is currently elusive.

In an attempt to reveal components that mediate the LINC-dependent arrest of DNA endoreplication, we performed a screen for genes whose transcription changes in the Nesprin/*klar* homozygous mutant muscles. One of the identified genes was *barrier-to-autointegration factor* (*baf*) (Wang et al., 2018). BAF is a small 89-aa protein, that forms dimmers, which are able to bind dsDNA, and thus capable of bridging between two strands of DNA, and thereby, to potentially affect DNA condensation (Bradley et al., 2005; Zheng et al., 2000). Alternatively, BAF dimers may bind to inner components of the nuclear membrane, including Lap-2, emerin, MAN1 (LEM) domain proteins, or to lamin A/C or B, as well as to dsDNA, and thus promote the association of dsDNA with the nuclear membrane (Cai et al., 2001; Lee et al., 2001; Mansharamani and Wilson, 2005; Samwer et al., 2017). Proteomic analysis of BAF partners indicated its potential association with additional proteins, including transcription factors, damage-specific DNA binding proteins, and histones (Holaska et al., 2003; Montes de Oca et al., 2009). BAF binding to its potential partners may be regulated by its phosphorylation state (Birendra et al., 2017; Lancaster et al., 2007; Nichols et al., 2006). For example, phosphorylated BAF has been shown to associate with LEM domain proteins, whereas de-phosphorylated BAF favors its binding to dsDNA (Bengtsson and Wilson, 2006; Nichols et al., 2006). This suggests that the various functions of BAF are controlled by specific kinases or phosphatases, or both. One kinase that has been implicated in BAF phosphorylation is VRK1 kinase, whose homolog in *Drosophila* is Ballchen (Ball also known as NHK-1) (Lancaster et al., 2007).

BAF has a crucial role in the condensation and assembly of post-mitotic DNA. Its interaction with both dsDNA and the nuclear lamina enables DNA compaction through cross-bridges between chromosomes and the nuclear membrane, a process that is essential for the formation of a single nucleus following mitosis (Samwer et al., 2017). Likewise, BAF is recruited to sites of ruptured nuclear membrane, where it is essential for sealing of these ruptures (Halfmann et al., 2019). Interestingly, in humans a single amino acid substitution of BAF causes Nestor–Guillermo progeria syndrome (NGPS) (Puente et al., 2011); however, the molecular basis for the disease awaits further investigation.

In *Drosophila* muscles, BAF transcription was reported to decrease significantly in mutant larvae lacking *Nesprin/Klar* or *SUN/koi*. Furthermore, muscle-specific knock down of BAF phenocopied the increased DNA endoreplication observed in LINC mutant myonuclei (Wang et al., 2018). This led to the hypothesis that BAF acts downstream of the LINC complex-mediated mechanotransduction, to arrest DNA endoreplication in muscles.

Here, we demonstrate that in mutants *Drosophila* larvae lacking functional LINC complex, BAF protein specifically dissociates from the nuclear membrane. Knockdown of BAF led to increased nuclear levels of E2F1 and Yap/Yorkie, two key positive regulators of cell cycle. Our evidence indicates that BAF forms a protein complex with E2F1, and that this interaction depends on BAF phosphorylation. Further, our results show that BAF kinase, VRK1/Ballchen, localizes at the muscle nuclear membrane, and its expression regulates BAF localization at the same site. Our findings suggest a model in which mechanosensitive localization of BAF at the nuclear membrane negatively regulates cell cycle progression by inhibiting accumulation of E2F1 and YAP/Yorkie in the myonuclei.

## Results

### BAF localization at the nuclear membrane depends on functional LINC complex

To test the hypothesis that BAF is involved in a mechanotransduction pathway that regulates cell cycle progression, we first analyzed the distribution of BAF protein in wild-type *Drosophila* larva muscles and then compared it to that of LINC mutant larvae. BAF was detected at the nuclear membrane (Fig 1 A-A’’), around the nucleolus (Fig 1 A, F), as well as in the cytoplasm (Fig 1 A, A’’). Notably, the labeling along the nuclear membrane overlapped that of lamin C, however it exhibited a relatively wider expression profile (Fig 1 A’’ and D). Furthermore, at the nuclear membrane BAF partially overlapped the labeling of *α*-tubulin (Fig 1 E), and since the latter is localized at the outer nuclear membrane (Elhanany-Tamir et al., 2012), this suggested that BAF localizes at both sides of the nuclear membrane. To determine whether BAF protein localization changes in the LINC mutants, we stained larvae homozygous mutant either for *koi,* or double homozygous for *klar* and *Msp30*0 alleles lacking only the KASH domain (*klar^ΔKASH^;Msp300^ΔKASH^*), which together represent the entire repertoire of LINC genes in the *Drosophila* genome. In both LINC mutants, BAF localization was specifically eliminated from the nuclear membrane, whereas its distribution around the nucleolus, and in the cytoplasm appeared similar to that of control (Fig 1 B-C’’). Automated, unbiased quantification of BAF-positive fluorescent voxels within the entire nuclear volume defined by lamin C outlines (see Materials and Methods) indicated a statistically significant reduction in the fluorescence levels of nuclear BAF, in muscles of both LINC mutants (Fig 1G). For quantification of BAF fluorescent levels, 66 myonuclei of control, 136 myonuclei of *koi* mutants, and 74 myonuclei of *klar^ΔKASH^;Msp300^ΔKASH^* mutants (in each group 4 muscles no. 7, from 4 distinct larvae) were analyzed. *t* test for *koi* indicated a significant reduction of nuclear BAF relative to control with p value <0.0001. Likewise, *t* test for *klar^ΔKASH^;Msp300^ΔKASH^* indicated a significant reduction relative to control with p value <0.0001. Larvae of all groups were staged, and labeled in parallel with the same antibody mix. These results implied that the localization of BAF protein at the nuclear membrane specifically depends on a functional LINC complex.

**Figure 1:**
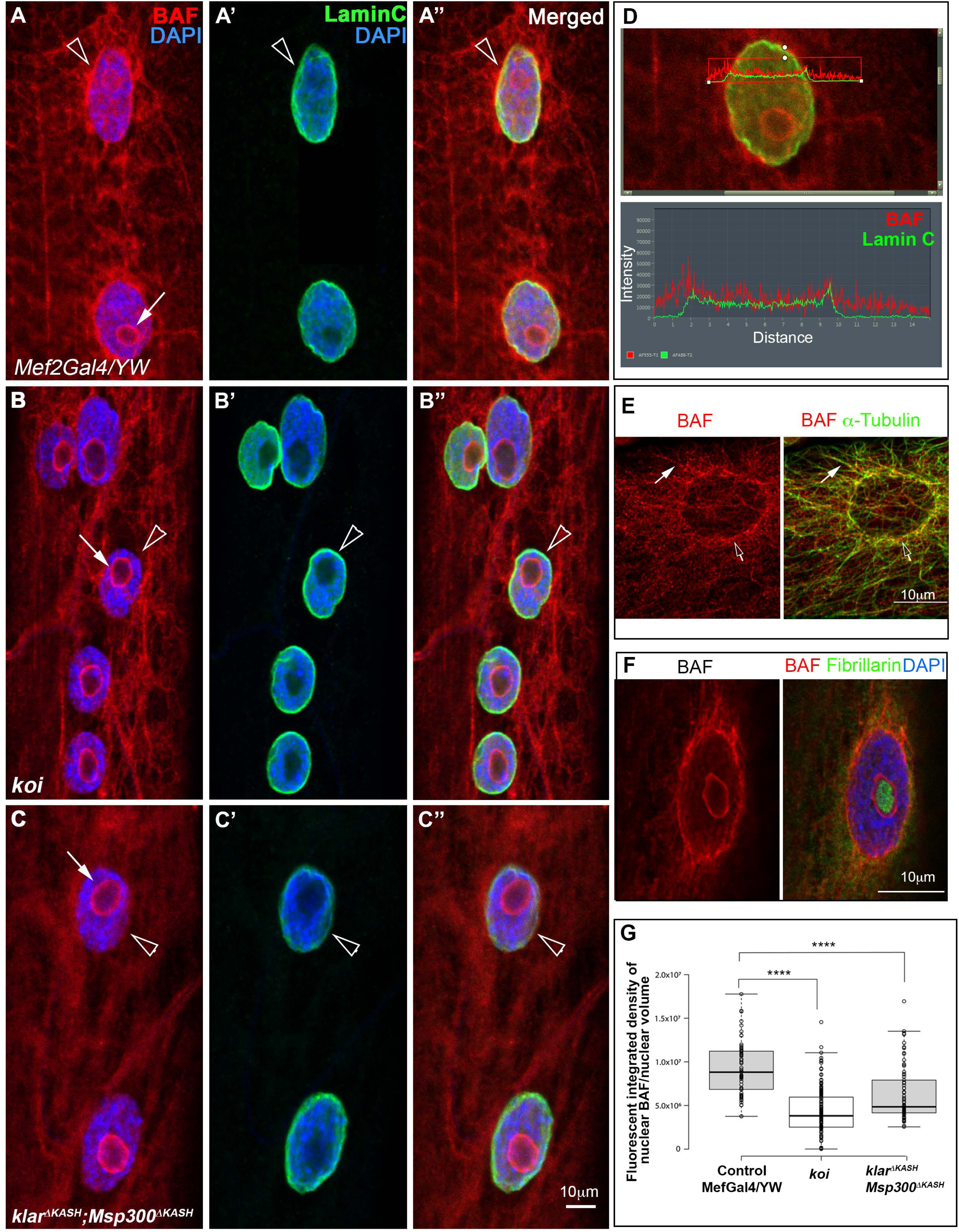
BAF dissociates from the nuclear membrane in LINC mutant muscles. A-C’’) Third instar larval muscle (no. 7) from control (Mef2-Gal4/YW, A-A’’) or LINC mutants, including either *klaroid/SUN* (*koi)* (B-B’’), or a double mutant combination of *klar^ΔKASH^* and Msp*^Δ^*^KASH^, in which the KASH domain from the two *Drosophila* Nesprins, *klar* and *Msp300* (C-C’’) was deleted, labeled with anti-BAF (red, A,B,C), lamin C (green A’,B’,C’) and their merged images (A’’,B’’,C’’). DAPI labeling is shown in blue (A,B, C). Arrows indicate the nucleolus, arrowheads indicate the nuclear membrane. D) upper panel: expression domains of BAF (red) and lamin C (green), (upper panel), and their relative signal intensities (lower panel) show that BAF is more widely distributed. E) A single myonucleus labeled with anti-BAF (red) and anti-*α*-tubulin (green), indicating partial overlap between BAF and tubulin (white arrows indicate overlapping labeling along microtubules, and empty arrows indicate non-overlapping labeling of BAF). F) A single myonucleus labeled with anti-BAF (red) and nucleolus marker Fibrillarin (green). Note that BAF labeling surrounds but not overlaps that of Fibrillarin. All images represent single confocal Z stacks. G) Quantification of the fluorescence integrated density of BAF per nucleus in control, *koi* mutant, or *klar^ΔKASH^*;Msp*^Δ^*^KASH^ double mutant myonuclei. *t* test for each of the LINC mutants relative to control indicates p<0.0001 (****). Bars in all images indicate 10μm.

Since we showed previously that BAF transcription is reduced in *koi*, and *klar* LINC mutants (Wang et al., 2018), we attempted to rescue BAF localization by its muscle-specific overexpression in *koi* mutant muscles. However, despite an increase in its cytoplasmic levels, BAF overexpression in *koi* mutants did not rescue its localization at the nuclear membrane (Fig 2 A-B’’).

**Figure 2:**
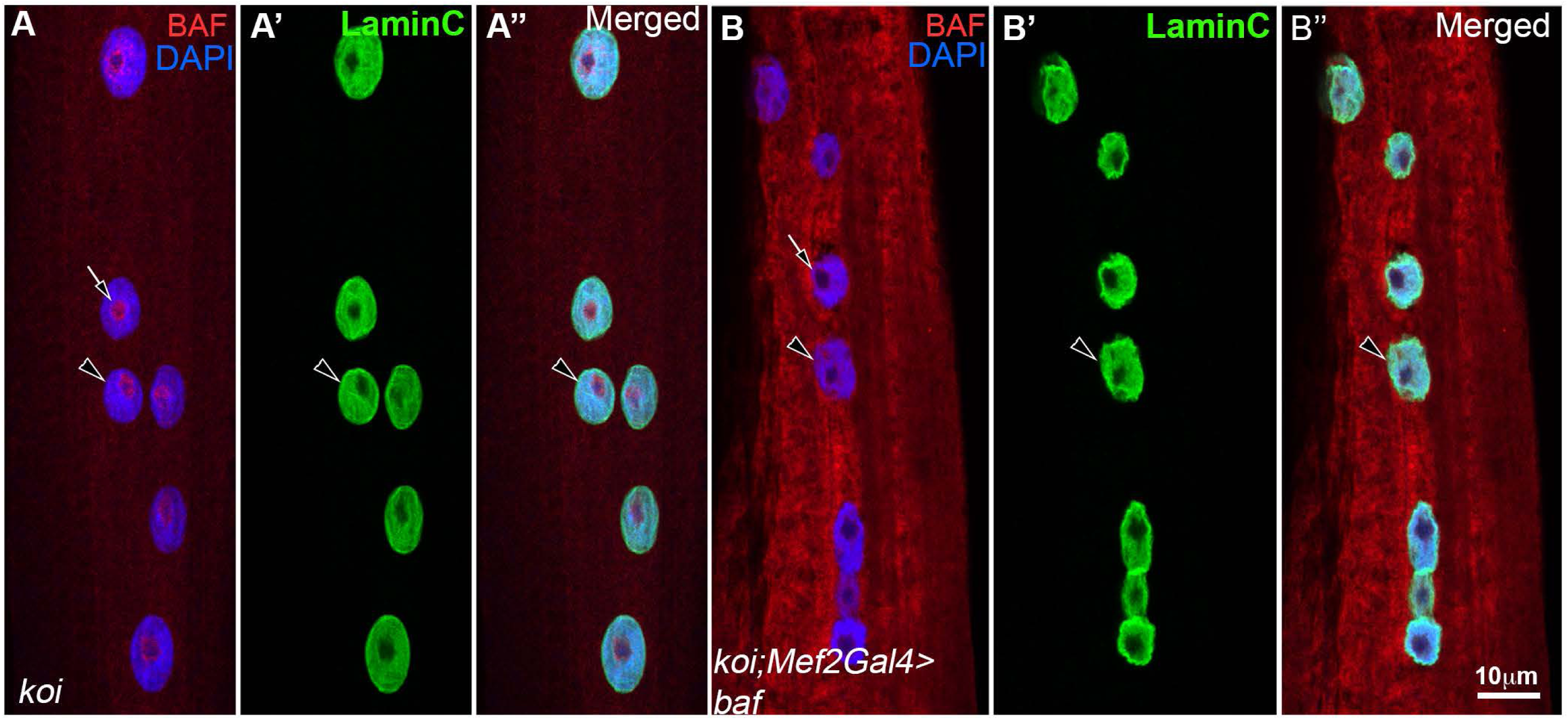
Overexpression of BAF in SUN/koi mutants does not rescue BAF at the nuclear membrane. A-A’’) larval muscle no. 7 from homozygous *SUN*/*koi* mutant, or from a *SUN/koi* mutant muscle overexpressing BAF (UAS-BAF) under the control of Mef2Gal4 driver (B-B’’), labeled with anti-BAF (red, A, A’’, B, B’’), anti-lamin C (green, A’, A’’, B’, B’’), DAPI (blue, A, A’’, B, B’’) and their merged images (A’’, B’’). Arrows indicate BAF at the nucleolus, and arrowheads indicate the nuclear membrane, which does not stain for BAF. All images represent a single confocal Z stack. Bar indicates 10 μm.

Taken together, these results show that BAF protein localization at the nuclear membrane depends on a functional LINC complex.

### BAF localization at the nuclear membrane depends on proper connection between myonuclei and sarcomeres

Previous studies indicated that myonuclei associate with the muscle sarcomeres through a functional interaction between D-Titin (Sallimus or Sls) and Msp300 (Elhanany-Tamir et al., 2012). To address whether myonucleus-sarcomere connections are required to maintain BAF at the nuclear membrane, a temporal, muscle-specific knockdown of *D-Titin/Sls* was induced at third instar larval stage using *sls* RNAi. Results show that in mutant muscles the myonuclei partially detached from the sarcomeres and they acquired rounded shape, in contrast to the flattened appearance of control myonuclei, implying altered mechanical input applied on the mutant nuclear membrane (Figure 3 E,F). Importantly, in the mutant myonuclei BAF dissociated from the nuclear membrane, whereas its localization around the nucleolus remained similar to that of control (Fig 3 A-B’’, and quantified in C). Quantification shown in Fig 3 C was based on 72 myonuclei (control) and 66 myonuclei of larvae expressing *D-Titin*/*sls-RNAi* (taken from 4 distinct muscles no. 7, and from 4 distinct larvae). A significant reduction in BAF levels at the nuclear membrane was observed for *D-Titin*/*sls-RNAi* expressing group relative to control, with *t* test indicating p value <0.0001. BAF at the nuclear membrane was determined by lamin C borders, and measured exclusively at this site (see Materials and Methods). These results indicated that BAF localization at the nuclear membrane depends on proper connection between the myonuclei and sarcomeres, and is sensitive to nuclear deformations arises following disruption of these connections. Notably, the DNA content in the myonuclei of the larvae expressing *D-Titin/sls* RNAi, quantified by DAPI fluorescent integrated density, increased significantly relative to control (Fig 3 D). DAPI fluorescence integrated density quantification was based on 67 myonuclei of control and 61 myonuclei of *D-Titin/sls* RNAi, each analyzed from 4 distinct muscles (no. 7), taken from 4 distinct larvae. The difference between the groups was significant, designated by *t* test with p value < 0.0001. Thus, reduced BAF levels at the nuclear membrane (induced by *D-Titin/sls* RNAi) correlated with an increase in DNA content (induced by elevated endoreplication) in these myonuclei. Previously, we reported that reduced BAF levels in muscles led to increased DNA levels in the myonuclei (Wang et al., 2018), whereas here we report that specific reduction of BAF at the nuclear membrane is sufficient for promoting increased DNA replication. Taken together these results demonstrate that BAF localization at the nuclear membrane is mechanosensitive, and might repress DNA endoreplication.

**Figure 3:**
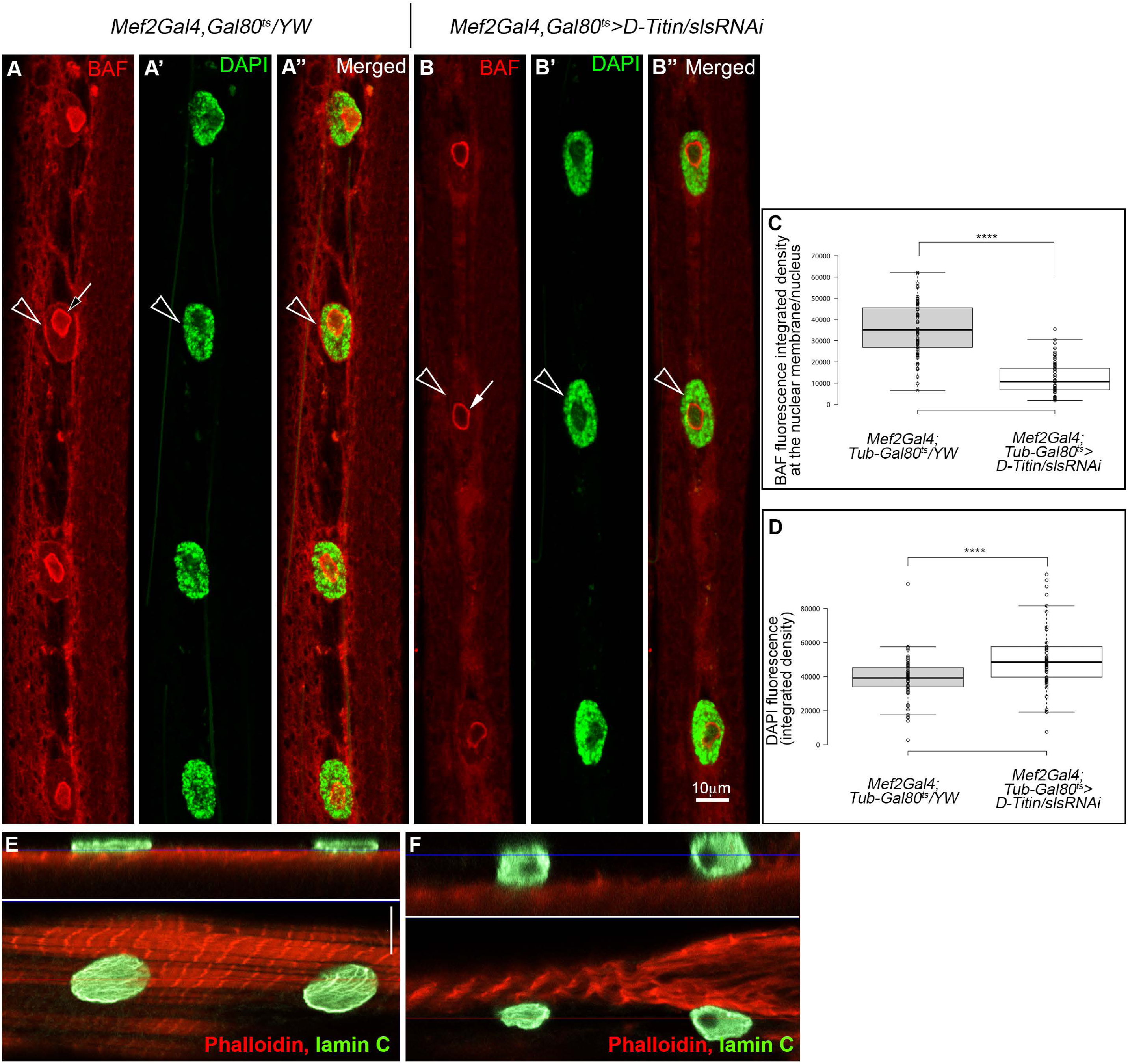
Partial dissociation of nuclear-sarcomeres connection by temporal knockdown of D-Titin/sls eliminates BAF from the nuclear membrane. A-B’’) larval muscles (no. 7) from control (*Mef2Gal4, Gal80^ts^* A-A’’, and E), or larvae temporally expressing *D-titin/sls* RNAi (*Mef2Gal4, Gal80^ts^*>*D-Titin/sls* (B-B’’, and F), grown in permissive temperature (18°C), and shifted to restrictive temperature (29°C) at second instar larval stage. A-B’’) larvae were labeled with anti-BAF (red, A, A’’, B, B’’) and with DAPI (green, A’, A’’, B’, B’’). Arrows indicate BAF at the nucleolus, and arrowheads indicate BAF at the nuclear membrane. C,D) Quantification of the fluorescence integrated density of BAF at the nuclear membrane per nucleus in control and in larvae temporally expressing *D-titin/sls* RNAi. BAF was significantly reduced at the nuclear membrane of *D-Titin/sls* knockdown muscles (C), whereas DAPI levels increased (D). E,F) Orthogonal (upper panels) and single X-Y optical section (lower panels) of muscle (no. 7) of control (E, Mef2Gal4, Gal80^ts^/YW) or *D-Titin/sls* knockdown muscles (F, Mef2Gal4, Gal80^ts^>*D-Titin/sls*), labeled with phalloidin (red, sarcomeres), and anti-lamin C (green). Red line indicates the location of the cross section, and blue line indicates the location of the X-Y optical section. Images (A-A’’) represent a single confocal Z stack. Bars indicate 10μm.

### E2F1 levels anticorrelate with BAF levels in the nucleus

To reveal the mechanism underlining the inhibitory effect of BAF on DNA replication, we focused on E2F1, a key transcription factor in promoting cell cycle progression and DNA endoreplication (Zielke et al., 2011). We analyzed the levels of E2F1 in myonuclei in which BAF levels were reduced by RNAi. The *baf* RNAi line that we used led to reduction in the protein levels of BAF, when driving it with mef2-Gal4 driver (deduced from immunolabeling as shown in Supp Fig S1 A-B’). Using pan induction of *baf* RNAi with *arm*-Gal4 and harvesting third instar larvae, revealed a 5 folds reduction of *baf* mRNA levels by qPCR analysis (Supp Fig S1 C).

Labeling of E2F1 protein in wild type larval muscles indicated an accumulation of the protein at the myonuclear membrane, as well as within the nucleoplasm (Fig 4A-A’’). Notably, E2F1 levels were significantly increased in the nucleoplasm of larvae expressing *baf* RNAi (Fig 4 A-B’’ and C). Quantification of E2F1 fluorescence integrated density was based on 150 control myonuclei, and 106 myonuclei of *baf-* RNAi expressing larvae (analyzed in both groups from 4 distinct muscles no. 7, which were examined from 4 distinct larvae). A significant elevation of E2F1 nuclear levels was observed in the *baf* RNAi expressing larvae indicated by *t* test with p value <0.0001. DAPI integrated fluorescent density quantified for myonuclei of the two groups (described above), indicated a significant increase of DNA levels in the group of larvae expressing *baf*-RNAi (Fig 4 D), with *t* test indicating p<0.0001, and consistent with our previous report (Wang et al., 2018). These results suggest an inhibitory function for BAF on E2F1 nuclear accumulation, and suggest that the upregulation of DNA replication in BAF knockdown myonuclei resulted from higher nuclear levels of E2F1. To address whether BAF and E2F1 distributions overlap at the nuclear membrane, we further analyzed the distribution of these proteins using expansion microscopy (Chen et al., 2015; Jiang et al., 2018). This procedure allows a roughly 4-fold increase in tissue size, while preserving its 3D constituents. Labeling with anti-BAF, lamin C and E2F1, using expansion microscopy indicated a complete overlap between the localization of the three proteins at the nuclear membrane (Fig 4 E-E’’’). However, BAF labeling at the outer nuclear membrane and at the nucleolus borders was lost, possibly due to the procedure used for expansion microscopy. Taken together, these results are consistent with an inhibitory interaction between BAF and E2F1, which might occur at the level of the nuclear membrane, where both proteins overlap.

**Figure 4:**
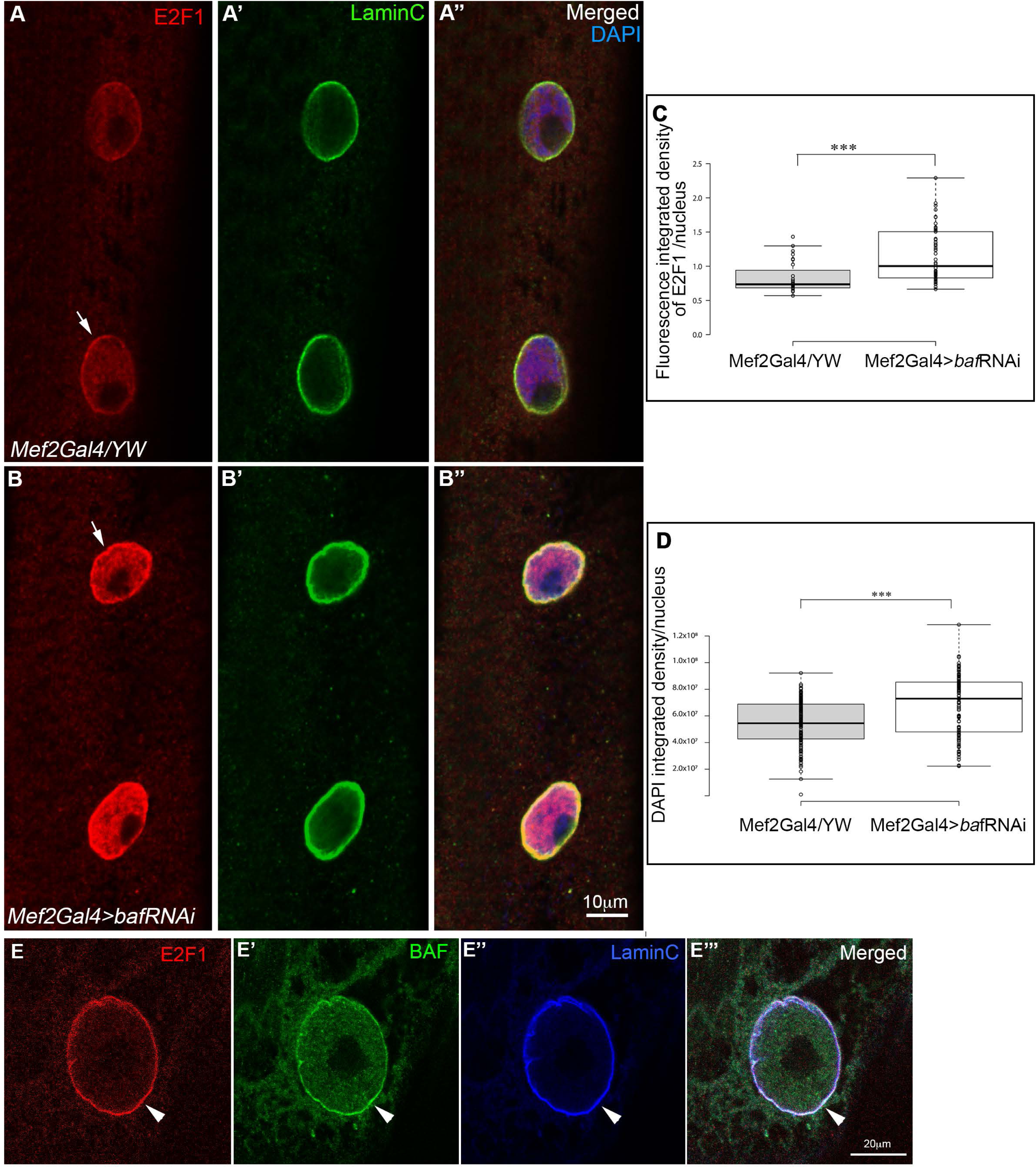
E2F1 nuclear levels increase following knockdown of BAF in muscles. A-B’’) Larval muscles (no. 7) of either control (Mef2Gal4/YW, A-A’’), or larvae expressing *baf* RNAi in muscles (*Mef2Gal4>baf RNAi*, B-B’’) labeled with anti-E2F1 (red, A, B), anti-lamin C (green, A’, B’) or their merged images (A’’, B’’) are shown. Arrows indicate the nuclear membrane. C) Quantification of the fluorescence integrated density of E2F1 per nucleus in myonuclei of control or BAF knockdown muscles. *t-*test indicates p< 0.0001 (****). D) Quantification of the fluorescence integrated density of DAPI per nucleus of myonuclei in control, or BAF knockdown muscles. *t-*test indicates p< 0.0001 (****). E-E’’’) single myonucleus using expansion microscopy indicates a complete overlap among E2F1 (red, E), BAF (green, E’), and lamin C (blue, E’’), as seen in their merged image (E’’’). Images in all panels represent a single confocal Z stack. Bars indicate 10μm or 20 μm.

### E2F1 and BAF form a protein complex

To test the hypothesis that BAF and E2F1 interact by forming a protein complex, we overexpressed GFP-BAF specifically in muscles. GFP-BAF was detected both in the cytoplasm and at the nuclear membrane, however, no enrichment at the nucleolus borders was observed (Fig 5A’). In these larvae E2F1 labeling was mainly detected at the nuclear membrane, overlapping that of GFP-BAF. BAF undergoes serine-threonine phosphorylation at three sites at its N-terminal end (see scheme in Fig 5) (Hou et al., 2016; Nichols et al., 2006). Previous studies indicated that non-phosphorylated BAF binds dsDNA, whereas phosphorylated BAF binds to LEM domain proteins at the nuclear membrane (Bengtsson and Wilson, 2006; Nichols et al., 2006). Thus, to examine the contribution of BAF phosphorylation to its inhibitory effect on E2F1, we performed muscle-specific overexpression of non-phosphorylatable BAF in which the three amino acids serine-threonine-serine at its N-terminal were replaced by 3 alanines (GFP-BAF-3A). Unfortunately, these larvae were significantly smaller and their muscles were severely disrupted. To overcome this problem, we induced temporal expression of GFP-BAF-3A in third instar larvae, as was described above for *sls*-RNAi). Examination of the larvae in these conditions indicated that the larvae grew up to a comparable size as control. Immunostaining revealed that GFP-BAF-3A accumulated specifically inside the nucleoplasm, overlapping DAPI and devoid of the nuclear membrane (Fig 5 B’). Notably, the nuclear levels of E2F1 in these myonuclei increased significantly relative to myonuclei overexpression GFP-BAF, grown in the same conditions (Fig 5 A, B and C). Quantification of E2F1 nuclear levels was based on 94 myonuclei of larvae expressing GFP-BAF, and 129 myonuclei of larvae expressing GFP-BAF-3A (analyzed from 4 distinct muscle no. 7, taken from 4 distinct larvae) in each group. A significant difference between the groups was observed by *t* test indicating p<0.0001 (Fig 5 C). Consistent with elevated levels of E2F1, the DAPI levels in the myonuclei overexpressing GFP-BAF-3A were significantly higher (Fig 5 D). Quantification of DAPI integrated density from these myonuclei (as above) showed a significant difference between the groups with *t* test indicating p<0.0001. These results suggested that excess of the nuclear GFP-BAF-3A, but not that of GFP-BAF released the BAF-dependent repression of E2F1 nuclear accumulation. We then used the GFP-BAF constructs for immunoprecipitation experiments to address whether E2F1 co-precipitates with BAF. Towards that end, larvae expressing either GFP-BAF, or GFP-BAF-3A, or control larvae (yellow-white, YW), all expressing the Mef2-GAL4, Tub-Gal80^ts^ constructs, were staged, and then grown in permissive temperature (18°C), and then transferred to restrictive temperature (29°C) at first instar larval for further growth. At third instar stage the larvae were dissected, and their protein extract was reacted with beads conjugated with anti-GFP antibodies for immunoprecipitation. The precipitated proteins were then reacted with anti-E2F1, as well as with anti-GFP. This analysis indicated that E2F1 specifically and significantly co-precipitated with wild-type GFP-BAF, but not with the non-phosphorylatable GFP-BAF-3A (Fig 5 E, F). It was therefore concluded that BAF and E2F1 form a protein complex, which does not form when BAF is not phosphorylated.

**Figure 5:**
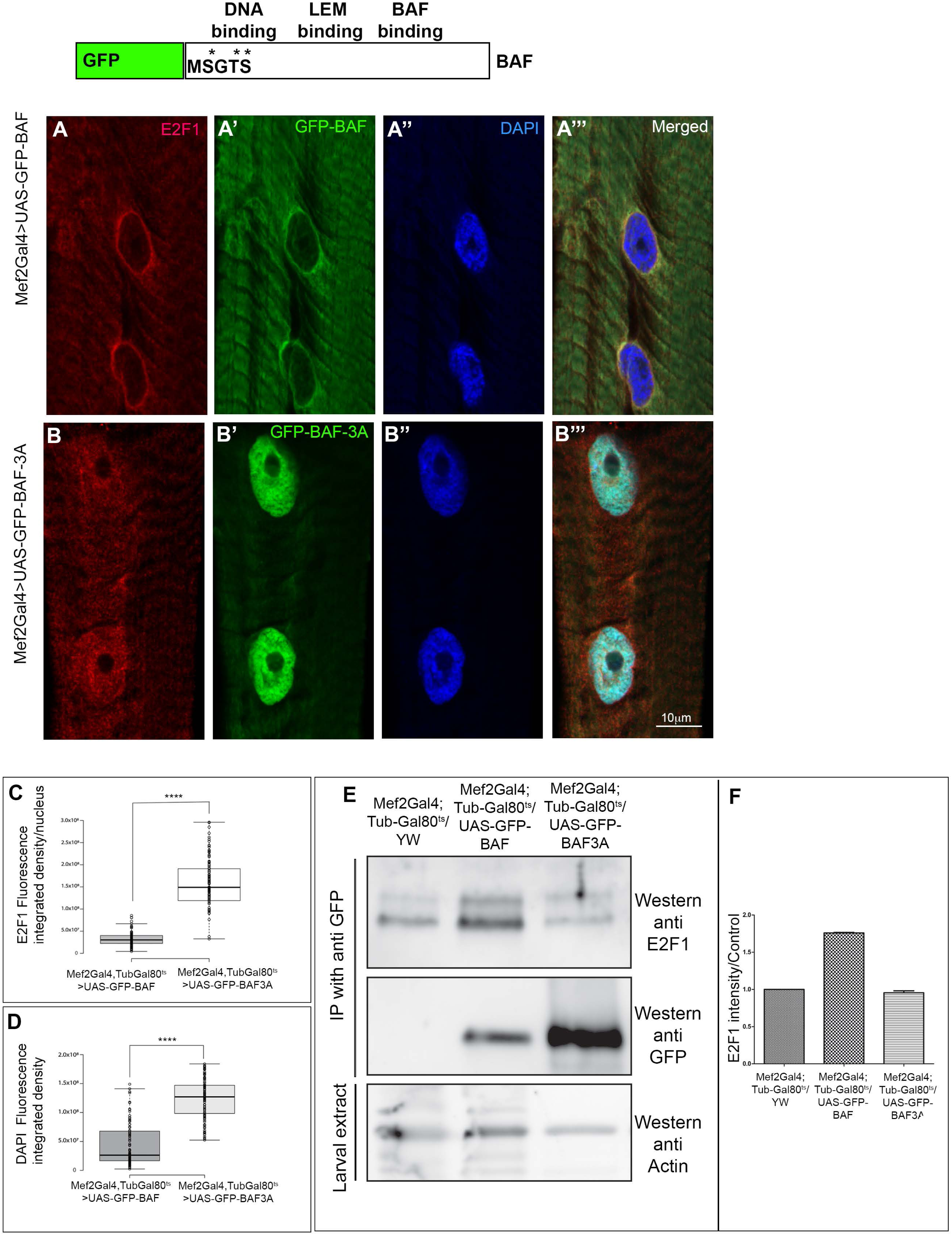
GFP-BAF co-precipitates with E2F1. A-B’’’) Larvae overexpressing either GFP-BAF (A-A’’’), or GFP-BAF-3A (B-B’’’) in muscles and labeled with anti-E2F1 (red, A, B, A’’’, B’’’), anti-GFP (green, A’, B’, A’’’, B’’’), and DAPI (blue, A’’, B’’, A’’’, B’’’) are shown. GFP-BAF is detected in both the cytoplasm and nuclear membrane, whereas GFP-BAF-3A is detected primarily in the nucleoplasm overlapping DAPI. Images in all panels represent a single confocal Z stack. Bar indicates 10μm. C) Quantification of E2F1 fluorescence integrated density/nucleus indicated higher levels in the myonuclei overexpressing GFP-BAF-3A relative to myonuclei overexpressing GFP-BAF. *t* test indicates p<0.0001. D-Quantification of DAPI fluorescence integrated density indicated higher levels in the myonuclei overexpressing GFP-BAF-3A relative to myonuclei overexpressing GFP-BAF. *t* test indicates p<0.0001. E) Co-immunoprecipitation of GFP-BAF or GFP-BAF-3A from larval extracts with anti-GFP beads, followed by Western blotting with anti-E2F1 indicated that GFP-BAF but not GFP-BAF-3A was capable of co-precipitating with E2F1 in levels higher than control. F) Quantification of the E2F1 bands relative to control in two distinct immunoprecipitation experiments.

### The serine/threonine kinase Vrk1/Ballchen regulates BAF localization at the nuclear membrane

Previous reports indicated that BAF is phosphorylated by the nuclear serine/threonine kinase Ballchen (Ball), a homolog of vertebrates Vrk1 (Bengtsson and Wilson, 2006; Lancaster et al., 2007). To address whether Vrk1/Ball controls the localization of BAF at the nuclear membrane, we knocked down Vrk1/*ball* in muscles (using *ball* RNAi), and followed BAF nuclear localization. Indeed, in these larvae BAF was absent from the nuclear membrane (Fig 6 A,B and C), whereas its localization at the nucleolus borders was retained. Quantification in Fig 6 C was based on measuring BAF fluorescence integrated density at the nuclear membrane, defined by lamin C borders. The analysis included 118 control myonuclei, and 69 myonuclei of larvae expressing *ball* RNAi (taken from 4 distinct muscle no. 7, and analyzed from 4 distinct larvae) in each of the groups. The difference between the groups was significant with *t* test indicating p<0.0001. This implied that Vrk1/Ball is required for BAF localization at the nuclear membrane, probably by promoting its phosphorylation. Interestingly, myonuclei of the Vrk1/*ball* knock-down muscles were not spaced evenly as in control, phenocopying the nuclear phenotype of the LINC mutants. Notably, the levels of E2F1 in the nucleoplasm of Vrk1/*ball* knock-down larvae were significantly elevated (Fig 6 A’,B’ and D). Quantification in Fig 6 D was based on 100 myonuclei of control and 126 myonuclei of larvae expressing *ball* RNAi (taken from 4 distinct muscles no. 7, analyzed from 4 distinct larvae). The difference between the groups was significant with *t* test indicating p<0.0001. These results are consistent with an inhibitory effect of Vrk1/Ball on E2F1 nuclear levels. Interestingly, using a fly line with endogenous GFP insertion into the *ball* locus, which enabled following its subcellular distribution revealed that Ball accumulated externally to lamin C labeling, presumably at the outer nuclear membrane (Fig 6 E, E’). Thus, the BAF kinase, Vrk1/Ball regulates the association of BAF with the nuclear membrane, presumably controlling the extent of its binding to LEM proteins.

**Figure 6:**
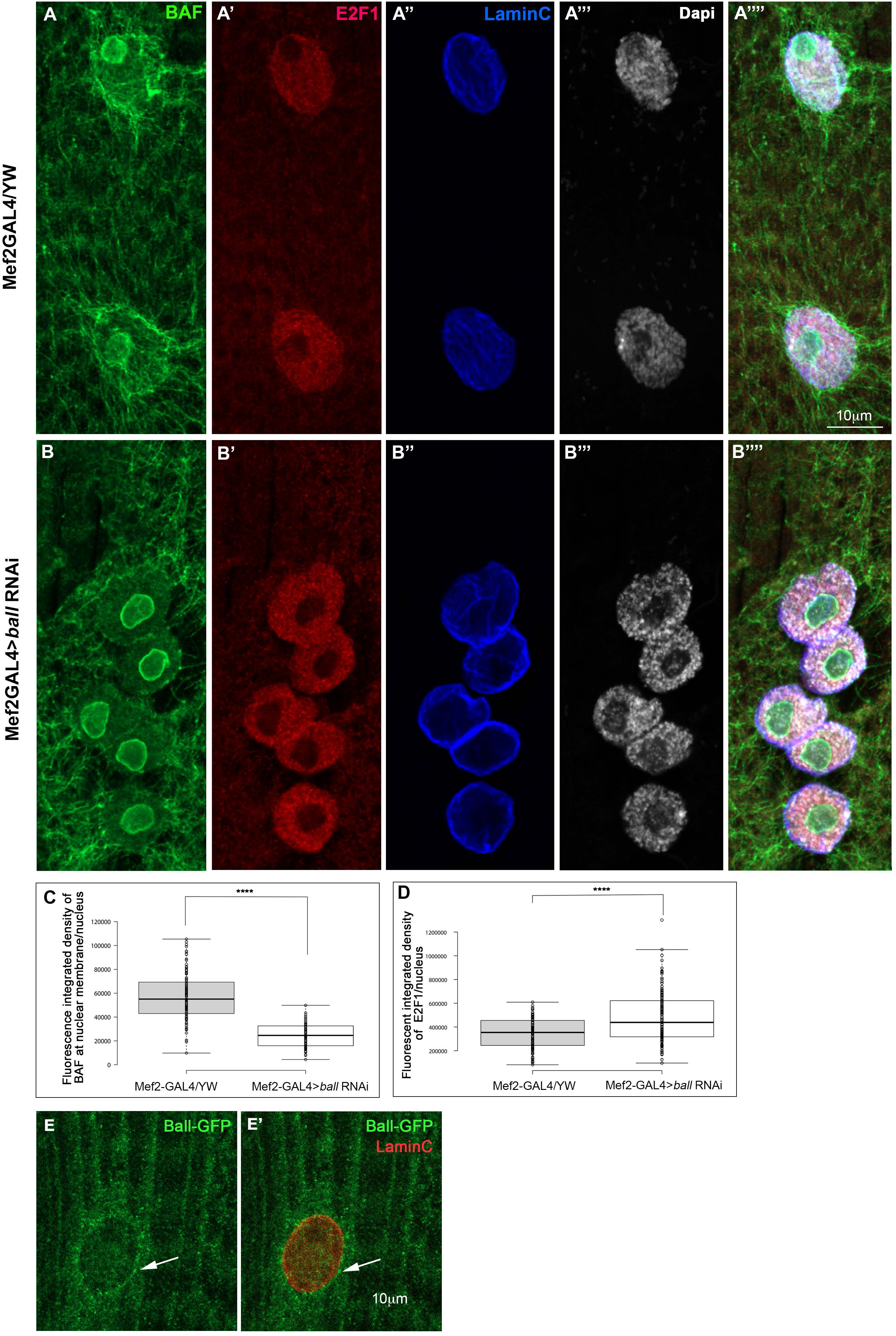
The Vrk1/Ball BAF kinase is required for BAF localization at the nuclear membrane. A-B’’’’) Larval muscles from control (Mef2Gal4/YW, A-A’’’’) or muscles expressing *Vrk1/ball* RNAi (Mef2Gal4>*Vrk1/ball RNAi*, B-B’’’’) were labeled with anti-BAF (green, A, B, A’’’’, B’’’’), anti-E2F1 (red, A’, B’ A’’’’, B’’’’), anti-lamin C (blue, A’’, B’’, A’’’’, B’’’’), and DAPI (white, A’’’, B’’’, A’’’’, B’’’’) and their merged images (A’’’’, B’’’’). A significant decrease in the levels of BAF at the nuclear membrane following knockdown of Vrk1/Ball was observed (A, B, C). C, D) Quantification of BAF fluorescence integrated density at the nuclear membrane (C) or E2F1 fluorescence integrated density in the nucleoplasm (D) indicated a significant decrease of BAF at the nuclear membrane (*t* test indicates p<0.0001), and a significant increase in nuclear E2F1 following knockdown of Vrk1/Ball (*t* test indicates p<0.0001). E, E’) Single myonucleus from larvae expressing GFP fused to endogenous *VRK1/ball*, labeled with anti-GFP (green, E,E’), and anti-lamin C (red, E’). Images in all panels represent a single confocal Z stack. Bar indicates 10 μm.

### BAF represses the accumulation of Yap/Yorkie in the myonuclei

Yap/Yorkie is an important mechanosensitive regulator of cell cycle (Dupont et al., 2011; Huang et al., 2005). To address whether in addition to E2F1, BAF also represses Yap/Yorkie, we used a YFP-tagged version of Yorkie expressed under its endogenous promoter (Yki-YFP), shown previously to rescue *yki* mutants (Su et al., 2017), to follow Yorkie subcellular distribution in muscles. Staining of control larval muscles with anti GFP antibody (which recognizes YFP), indicated homogenous distribution of Yki-YFP within the muscle with no accumulation in the myonuclei. However, muscles in which BAF was knocked down with RNAi, exhibited a significant accumulation of Yki-YFP in the larval myonuclei (Fig 7 A-C). Quantification of nuclear Yki-YFP was based on 66 myonuclei of control larvae, carrying YFP-Yorkie and Mef2GAL4 (taken from 9 muscles no. 7 analyzed from 3 larvae), and 73 myonuclei from larvae expressing in addition *baf* RNAi (taken from 9 muscles no. 7 from 4 larvae). The difference between the groups was significant with *t* test indicating p<0.0002. This result suggests that BAF represses the nuclear entry of Yap/Yki.

**Figure 7:**
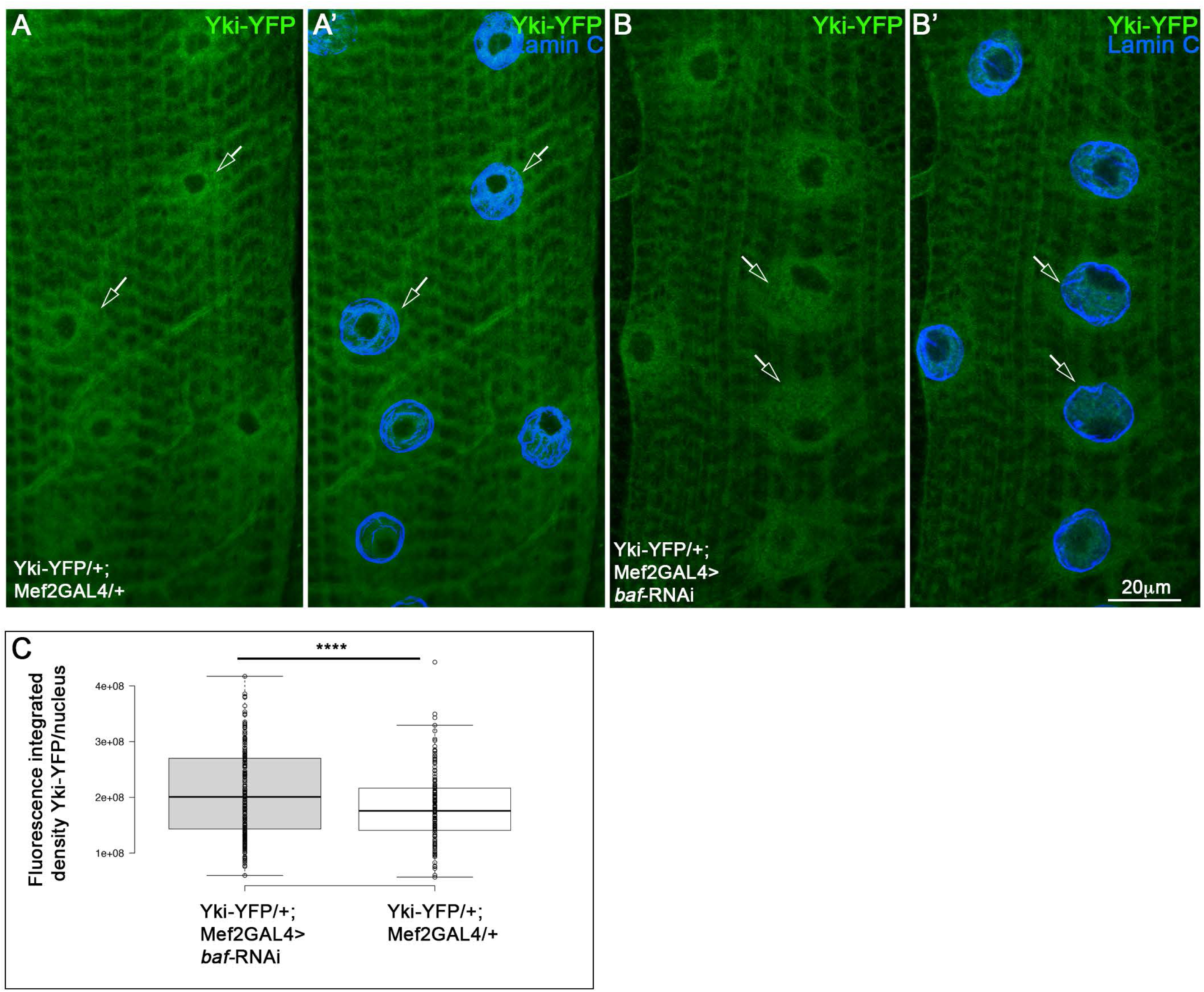
Yap/Yorkie nuclear levels are elevated following BAF knock down in myonuclei. A-B’) Larval muscles carrying Yap/Yorkie-YFP under endogenous control and combined with either Mef2Gal4 (A,A’), or Mef2Gal4 with *baf* RNAi (B,B’) were labeled with anti-YFP (green) and anti-lamin C (blue). Arrows indicate the border of the nuclear membrane, labeled with lamin C. Images in all panels represent a single confocal Z stack. C) Quantification of the fluorescent integrated density in the myonuclei indicated a significant increase in the levels of nuclear Yap/Yki-GFP following knockdown of BAF (*t* test indicates p<0.0001). Bar indicates 20μm.

## Materials and methods

### Fly stocks and husbandry

All crosses were performed at 25°C and rose on cornmeal yeast agar. The following stocks were distributed by the Bloomington Drosophila Stock Center: tubP-GAL80^ts^/TM2 (FBst0007017), GAL4-Mef2.R (FBst0027390), *baf*-RNAi (FBst0036108), *sls* RNAi (FBgn0086906), *ball* RNAi (FBti0130755). Ball-GFP was obtained from Vienna BioCenter (VDRC): FlyFos016090 (pRedFlp-Hgr)(ball[41529]::2XTY1-SGFP-V5-preTEV-BLRP-3XFLAG)dFRT. UAS-GFP-BAF, UAS-GFP-BAF-3A (obtained from Lei Liu, Beijing Institute for Brain Disorders, Beijing, China). *klar^ΔKASH(mCD4)^ and Msp300^ΔKASH^* (J.A. Fischer, University of Texas, Austin, TX) were combined on Cyo-YFP and TM6Tb balancers, *koi^84^*(FBst0025105)/Cyo-dfd-eYfp (Kracklauer et al., 2007). Yorkie-YFP (R. Fehon, University of Chicago (Su et al., 2017).

Staging of the larvae was performed by 6 hours embryo collection and further growing on yeast past in vials up to early third instar stage. Temporal expression of *sls* RNAi or GFP-BAF, as well as GFP-BAF-3A was performed by using a combination of Mef2GAL4, tubGal80^ts^ drivers, as follows: embryos collection was performed at 25°C for 6 hours, then embryos were transferred to permissive temperature 18°C up to first instar larval stage, and then larvae were transferred to restrictive temperature of 29°C, up to early third instar stage.

### Immunofluorescence

Immunofluorescent staining were performed as previously described (Wang et al., 2015). For fixation, paraformaldehyde (4% from 16% stock of EM grade; #15710; Electron Microscopy Sciences) was used without methanol to avoid damage of native F-actin or chromosomal morphology. Specimens were fixed for approximately 30 minutes and subsequently washed several times in PBS with 0.1% Triton X-100 (PBST) on a horizontal shaker with gentle agitation. Image analyses were consistently performed on muscle 7. All specimens were mounted in Thermo Scientific Shandon ImmuMount for microscopy (Thermo Fisher Scientific).

### Antibodies and synthetic dyes

Mouse anti-lamin C (DSHB, no. LC28.26-c) obtained from the Developmental Studies Hybridoma Bank, created by the NICHD of the NIH and maintained at The University of Iowa. Rat anti-Tubulin alpha (Bio-Rad MCA78G), Chicken anti-GFP (Abcam #13970), Rabbit anti-BAF (provided by Paul Fisher, Stony Brook, NY, and Ryszard Rzepecki, University of Wroclaw, Poland, (Furukawa et al., 2003), Rat anti-E2F1 was used for immunofluorescence and Guinea Pig anti-E2F1 was used for Western analysis. Both antibodies were provided by provided by Stefan Thor (University of Queensland, Australia), (Baumgardt et al., 2014), and Jonathan Benito-Sipos, (University of Madrid). Secondary antibodies used: Alexa Fluor 488, 555 and 647 conjugated secondary antibodies against Rat, Chick, Rabbit, and mouse were purchased from The Jackson Laboratory and Thermo Fisher Scientific.

For labeling of the chromatin we used DAPI (1 µg/ml; Sigma-Aldrich). For F-actin labeling we used TRITC-Phalloidin (SIGMA-ALDRICH P1951).

### Expansion microscopy

The procedure was essentially as described (Jiang et al., 2018). Briefly, larvae were fixed with 4% PFA in PBS, washed with PBST, blocked with 10% BSA in PBST, incubated with first antibody overnight, and with secondary antibodies (conjugated with either Atto 647, AF555, AF488) overnight. After wash with PBST, and PBS the larvae were incubated with 1mM MA-NHSin PBS for 1hour, washed with PBS, incubated with monomer solution (2M NaCl, 8.625% Sodium Acrylate, 2.5% Acrylamide, 0.15% Bisacrylamide, 10X PBS) for 45 min at 4°C, and then transferred to gelation solution (0.2% TEMED, 0.01% TEMPO, 95% monomer solution, and 0.2% APS), for 30 min at 37°C. The gel with the larvae was incubated with chitinase (1Unit/mL) in PBS pH 6.0, 4 days at 37°C. Three wash with PBS, were followed by the addition of collagenase (1mg/mL) in 1x HBSS (w/0.01 M CaCl_2_, and 0.01 M MgCl_2),_ and incubated at 37°C O.N. Washes with PBS were followed by addition of Proteinase K (8Units/mL) in digestion buffer at 37°C, for 1 Hour and additional washes with PBS. Hoechst was added for 10 min, washed and expansion was performed by the addition of ultrapure water for 30 min. The sample was then imaged under the confocal microscope.

### Microscopy and image analysis

Microscopic images were acquired at 23°C on confocal microscopes Zeiss LSM 800 with the following lenses: Zeiss C-Apochromat 40x/1.20 W Korr M27, 20x PlanApochromat 20/0.8. The microscopic samples were embedded with Coverslip High Precision 1.5H ± 5 μm (Marienfeld-Superior, Lauda-Königshofen, Germany). Immersion medium Immersol W 2010 (ne = 1.3339) and Immersion oil Immersol 518 F (ne = 1.518) were used, respectively. Images were analyzed with Fiji including plug-ins and adapted scripts. Figure panels were finally assembled using Photoshop CC 2019). Acquisition software Zen 2.3 (blue edition).

### Automatic Data Collection

Quantification of fluorescence integrated density of proteins in myonuclei was performed using a custom-built FIJI macro which included rolling ball background subtraction. The macro has been deposited in GitHub DOI 10.5281/zenodo.3372266. In brief, lamin C was used to define the entire nuclear volume as the region of interest (ROI). In each myofibril integrated fluorescent density in different channels were measured in all the slices along the Z-direction of the ROI. This allowed us to measure the levels of specific proteins inside the myonuclei in an automated and unbiased manner. For measuring BAF levels along the membrane, we used the manual selection tool for the lamin C channel to define the nuclear membrane as the ROI. The fluorescence integrated density of BAF along the Z-direction of the ROI was then analyzed.

### Statistical analysis

Statistical analysis was performed using SPSS software (IBM SPSS for Windows, Version 19) and Microsoft Excel 2013. Measurements were first evaluated for normality using the Shapiro-Wilk test, and subsequently analyzed using a two-independent samples t-test. P-values <0.05 were considered statistically significant. BoxPlotR was used to generate box plots (Spitzer et al., 2014), in which the center lines represent the medians, box limits indicate 25th and 75th percentiles, crosses represent sample means, and outliers (data points beyond the 95% confidence intervals) are represented by dots. Furthermore, whiskers, determined using the Tukey method, extended to data points less than 1.5 interquartile ranges from the 1st and 3rd quartiles as determined by the BoxPlotR software. All experiments were performed at least twice independently and in at least 4-5 randomly selected larvae from which muscle no. 7 was monitored in at least 4 abdominal segments, per experimental group.

#### Western blot and immunoprecipitation

Third instar larvae were collected and protein extract was prepared in modified RIPA buffer [1% NP40, 150 mM NaCl and 50 mM Tris pH 8.0] supplemented with protease inhibitor cocktail (Sigma). For immunoprecipitation, equal amounts of protein lysate were incubated overnight at 4°C with GFP-TrapR_A beads (Chromotek, BioNovus Life Sciences). Beads were boiled in SDS sample buffer, separated by SDS–PAGE and subjected to western blot analysis with the indicated antibodies. Primary antibodies used included chick anti-GFP (Abcam ab13970) Guinea Pig anti-E2F1. Secondary antibodies used were either anti-chick HRP or anti-Guinea Pig HRP (Jackson ImmunoResearch). Chemiluminescent detection was performed using SuperSignal West Pico (Thermo Scientific), according to the manufacturer’s directions.

#### RT-qPCR

Gene expression quantification was carried out by reverse transcription quantitative real time PCR (RT-qPCR). Briefly, total muscle RNA was isolated from larval body walls of 30 dissected Drosophila individuals using RNeasy Protect Mini Kit (Qiagen). 1 μg of total RNA were used for first strand cDNA synthesis by reverse transcription using SuperScript Reverse Transcriptase IV (Invitrogen) with oligonucleotide dT primers. qPCR was performed using gene specific primers on the ABI 7500 Real-Time PCR System (Applied Biosystems) with Fast SYBR Green Master Mix (Applied Biosystems) for detection. Each sample was run in triplicate. The primer sequences are listed in Table (see supplementary information Table 1). The difference in gene expression was calculated using the fold change (ΔΔCt method) (Schmittgen and Livak, 2008). ΔCt is the Ct value for the gene of interest normalized to the Ct value of the respective gene in both control larvae (armadillo-Gal4/YW) and BAF knockdown (armadillo-Gal4>*baf* RNAi). ΔΔCt values were calculated as a relative change in ΔCt of the target gene in BAF knockdown with respect to control - house-keeping gene *succinate dehydrogenase (SDH)*. Fold changes were expressed as 2− ΔΔCt for up-regulated genes and the negative reciprocal of the fold change for down-regulated genes (where 2− ΔΔCt < 1).

## Discussion

Mechanosensing by the nucleus has been implicated in a wide array of processes, including cell cycle progression (Cho et al., 2017; Kirby and Lammerding, 2018). Nevertheless, the molecular basis underlying mechanotransduction-dependent cell cycle regulation is not fully explained. Here, we demonstrate the contribution of a novel mechanosensitive component, BAF, in controlling the nuclear accumulation of two transcription factors, E2F1 and YAP/Yorkie, which have been implicated in regulation of cell cycle progression in a wide variety of cell types and tissues (Huang et al., 2005; van den Heuvel and Dyson, 2008; Zielke et al., 2011). Whereas previous reports implicated BAF in the condensation and assembly of post-mitotic dsDNA into single nuclei (Samwer et al., 2017), here we demonstrate that BAF is also essential for the arrest of DNA endoreplication in fully differentiated muscle fibers. Importantly, while this function was observed in post-mitotic differentiated cells, BAF may regulate the nuclear accumulation of E2F1 and YAP/Yorkie in dividing cells as well.

In the *Drosophila* muscle fibers, we found that BAF was detected in multiple subcellular sites, including the cytoplasm, the nuclear membrane, the nucleoplasm and surrounding the nucleolus. Yet, only the portion of BAF localized at the nuclear membrane was found to be sensitive to altered mechanical inputs, either in muscles lacking functional LINC complex, where myonuclei deform and become more rounded (Elhanany-Tamir et al., 2012), or in muscles expressing *sls* RNAi, in which myonuclei partially detached from the sarcomeres and undergo significant cell shape changes. In both scenarios, nuclear shape changes are indicative of altered nuclear mechanics induced by disruption of the cytoskeletal-nuclear connections, hence it was concluded that maintenance of BAF at the nuclear membrane is mechanically sensitive.

In control myofibers, BAF exhibited a relatively broad distribution along the outlines of the nuclear membrane, often extending beyond the lamin C expression domain towards the cytoplasm, where it was partially overlapped with the nuclear-associated microtubules. This suggests that in addition to its association with the inner aspects of the nuclear membrane through its binding to LEM-domain proteins and lamin A/C (Cai et al., 2001; Jamin and Wiebe, 2015; Shimi et al., 2004; Wilson et al., 2005), BAF also associates with the outer aspects of the nuclear membrane in myofibers. Interestingly, BAF localization on both sides of the nuclear membrane was disrupted following alterations in the mechanical effects on the nuclear membrane, implying that both inner and outer pools of BAF are sensitive to mechanical inputs and might be functionally connected. Previous experiments indicated that BAF does not diffuse passively from the cytoplasm to the nucleus (Shimi et al., 2004). Furthermore, photobleaching experiments with GFP-BAF indicated that BAF-dependent repair of nuclear ruptures occurs when cytoplasmic BAF, but not internal nuclear BAF, is recruited to the ruptured sites, and this further brings LEM domain proteins that are essential for membrane sealing (Halfmann et al., 2019). These findings are consistent with a dynamic exchange between cytoplasmic and nuclear BAF, in which BAF in the cytoplasm primarily responds to mechanical signals. Since we found that BAF kinase VRK1/Ball is predominantly localized at the cytoplasmic aspects of the nuclear membrane, it is possible that BAF phosphorylation is an early step in BAF response to mechanical signals.

It is not clear which portion of BAF becomes associated with E2F1. However, the distribution of E2F1 appeared more confined than that of BAF, and overlapped completely that of lamin C, supporting a model where the protein is retained at the inner aspects of the nuclear membrane together with BAF (see model in Fig 8). Since our Co-IP experiments suggest that non-phosphorytable BAF does not form a protein complex with E2F1, we suggest that BAF-E2F1 association depends on cytoplasmic BAF. Thus, we suggest that VRK1/Ball kinase phosphorylates BAF at the outer nuclear membrane and then pBAF translocated into the inner nuclear membrane, where it is tethered to the membrane by the LEM proteins. Such a scenario is consistent with a recently suggested model for BAF involvement in repair of mechanically induced membrane rupture (Halfmann et al., 2019). In both cases, recruitment of cytoplasmic BAF to the nuclear membrane is sensitive to changes in the mechanical properties of the nuclear membrane, and then, due to interaction with LEM proteins, BAF becomes localized to the inner nuclear membrane. Whether or not this sensitivity involves phosphorylation/dephosphorylation of BAF is yet to be determined.

**Figure 8:**
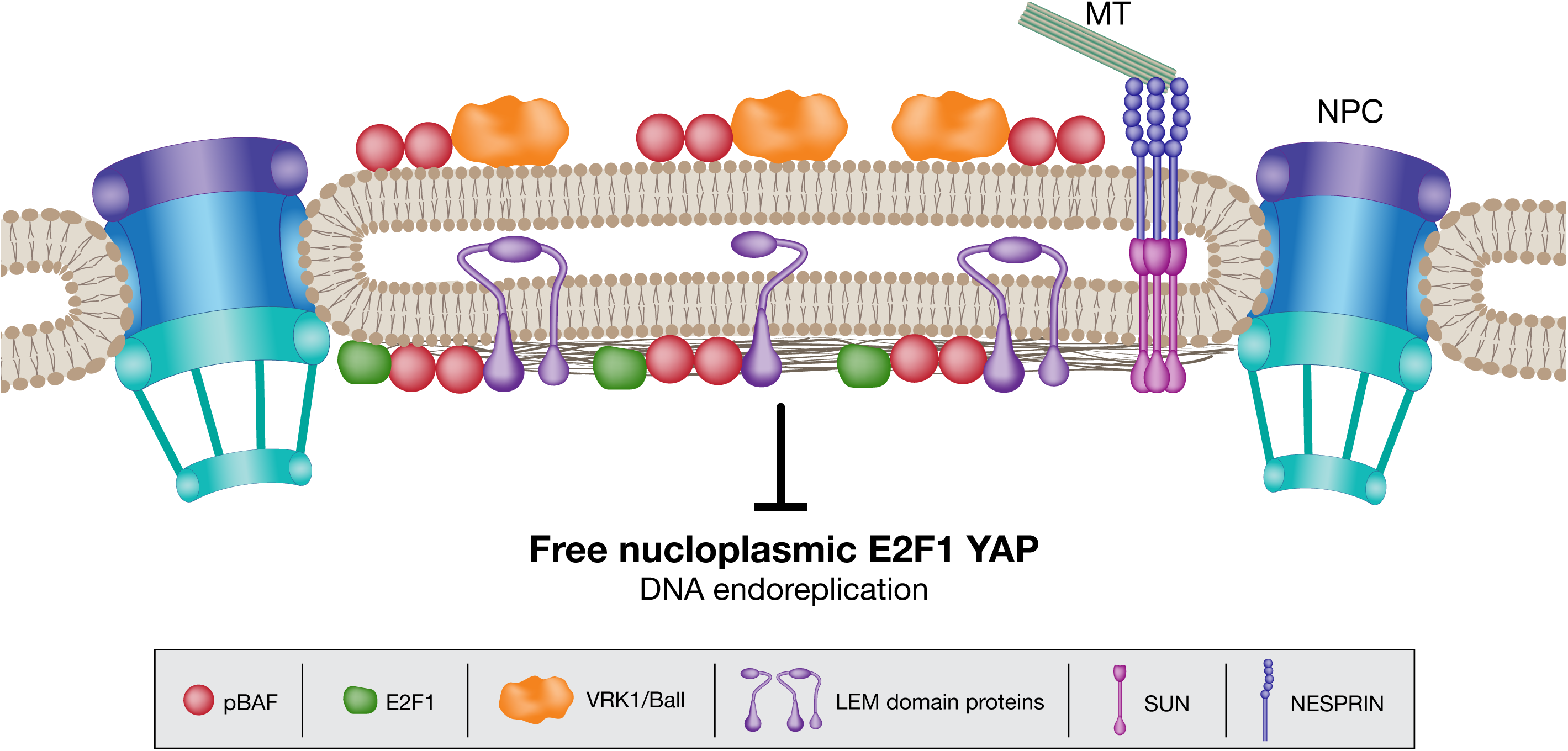
Model of BAF-dependent mechanosensitive repression of E2F1 and YAP/Yorkie. In the suggestive model, BAF association with the nuclear membrane results from its phosphorylation by VRK1/Ball in the presence of functional LINC complex. pBAF associate with E2F1 at the nuclear membrane, repressing its accumulation in the nucleoplasm, thereby inhibiting DNA endoreplication. YAP/Yorkie accumulation is also inhibited by BAF, but the mechanism of this inhibition is unclear.

Our results indicated that BAF is essential not only for E2F1, but also for Yap/Yorkie accumulation in the nucleoplasm. Previous studies showed that Yap/Yorkie nuclear translocation is sensitive to mechanical inputs (Aragona et al., 2013; Codelia et al., 2014; Cui et al., 2015; Driscoll et al., 2015; Sharili and Connelly, 2014). Furthermore, nuclear translocation of Yap/Yorkie was increased following application of mechanical forces on the nuclear membrane, and it required nuclear pore opening (Elosegui-Artola et al., 2017). It remains unclear whether the two processes, namely BAF phosphorylation and nuclear pore opening, are functionally related and hence represent a general, mechanically sensitive mechanism for nuclear translocation.

In summary, our results reveal novel insights into cell cycle regulation and show how it is linked to changes in the mechanical properties of the nuclear membrane in non-dividing cells. In particular, a novel mechano-sensitive component, BAF, is shown to negatively regulate the nuclear entry of key cell cycle regulators namely, E2F1 and Yap/Yorkie, at the level of the nuclear membrane. BAF phosphorylation by VRK1/Ball kinase is critical in this context, since it controls the association of BAF with the nuclear membrane. Importantly, this process may be part of a mechanosensitive pathway that regulates cell cycle progression in dividing cells.

## Acknowledgments

We thank Paul Fisher (State University of New York at Stony Brook), Ryszard Rzepecki (University of Wroclaw, Poland), Stefan Thor (University of Queensland, Australia) and Jonathan Benito-Sipos (University of Madrid) for providing antibodies, and Richard Fehon (University of Chicago), and Lei Liu (Beijing Institute for Brain Disorders) for providing fly lines. We are grateful for the Bloomington Stock Centre and Vienna Drosophila Resource Center (VDRC) for various fly lines, the Developmental Studies Hybridoma Bank (DSHB) for antibodies, and FlyBase for important genomic data. We are grateful to Ofra Golani (Weizmann Core facilities) for writing the script for quantification of fluorescent images, and also thank Nitzan Konstantin for English editing. This study was supported by grants from “The French Muscular Dystrophy Association (AFM-Téléthon)” grant # 22339, NSF-BSF (BSF grant # 2016738), and Israel Science Foundation (ISF) grant # 750/17.

---

**Supplementary Figure 1:**
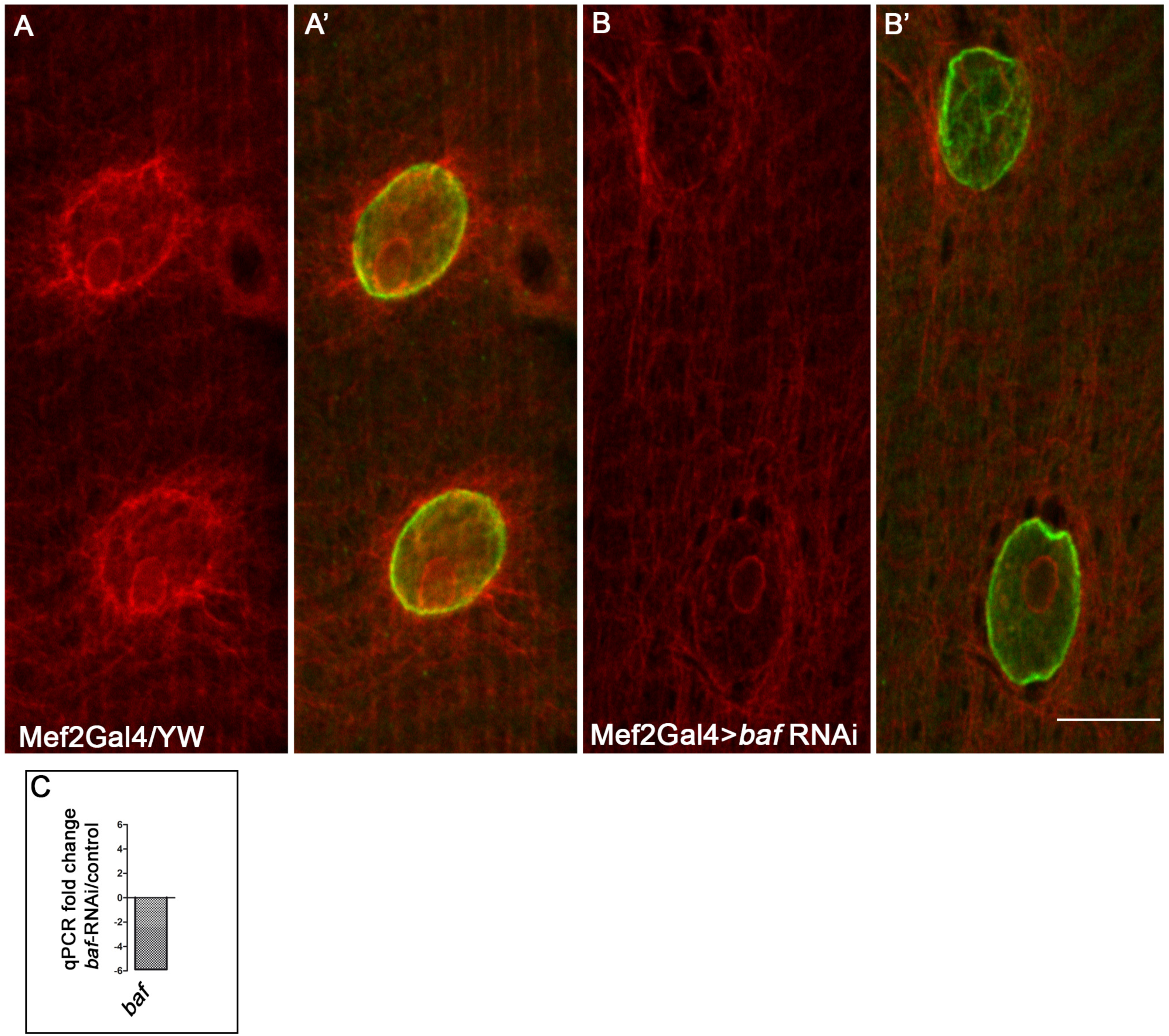
Evaluation of the efficiency of baf RNAi line. Larval muscles (no. 7) labeled with anti BAF (red, A-B’), as well as with lamin C (green A, B’) of control (Mef2Gal4/YW) or baf-RNAi (Mef2Gal4>baf-RNAi) larvae. Note a reduction in BAF protein levels. C) qPCR analysis using baf primers, as well as primers for house-keeping gene succinate dehydrogenase (SDH) (for normalization), of control larvae (armGal4/YW), or baf RNAi larvae (armGal4>baf-RNAi). This analysis indicated a 5 fold reduction in baf mRNA levels

