## Supplemental Figure 1 for "Mechanosensitive recruitment of BAF to the nuclear membrane inhibits nuclear E2F1 and Yap levels"

Supplementary Figure 1
.C.P. Unnikannan et al

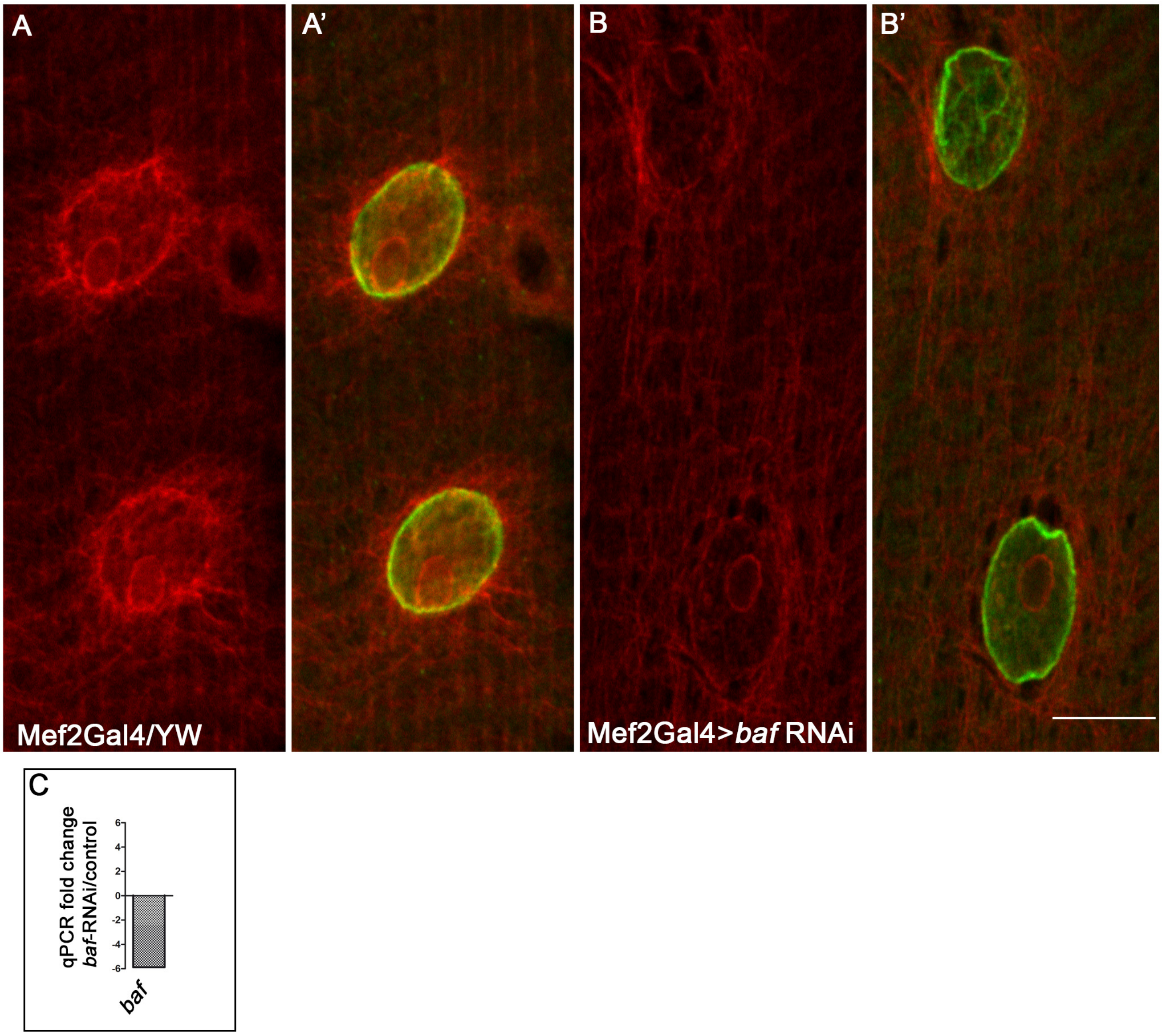


<u>Supplementary Figure 1: Evaluation of the efficiency of baf RNAi line</u>
Larval muscles (no. 7) labeled with anti BAF (red, A-B’), as well as with lamin C (green A, B’) of control (Mef2Gal4/YW) or baf-RNAi (Mef2Gal4>baf-RNAi) larvae. Note a reduction in BAF protein levels. C) qPCR analysis using baf primers, as well as primers for house-keeping gene succinate dehydrogenase (SDH) (for normalization), of control larvae (armGal4/YW), or baf RNAi larvae (armGal4>baf-RNAi). This analysis indicated a 5 fold reduction in baf mRNA levels
